# Mpox Vaccine Design Through Immunoinformatics and Computational Epitope Prediction

**DOI:** 10.1101/2024.11.05.622158

**Authors:** Sebastián Rivera-Orellana, José R. Ramírez‑Iglesias, Jaime David Acosta-España, Jorge Espinosa-Espinosa, Juan-Carlos Navarro, Andrés Herrera-Yela, Andrés López-Cortés

## Abstract

The Mpox virus (Monkeypox virus) poses significant public health risks due to its potential for severe outbreaks in humans. This study presents an innovative vaccine design using bioinformatics to identify epitopes that activate helper T cells (HTLs) via the human leukocyte antigen class II (HLA-II) complex. Starting with 50,040 vaccine candidates, 14 epitopes with the highest HLA-II affinity were selected based on antigenicity, allergenicity, toxicity, stability, and homology. These epitopes were integrated into a multi-epitope vaccine with spacers and adjuvants to enhance the immune response. A 3D model was developed, confirming structural stability and optimal epitope exposure through molecular dynamics simulations. The results indicate that the vaccine can induce robust immune responses, suggesting its potential effectiveness against the Mpox virus. Additionally, population coverage analysis supports its promise as a significant tool for controlling Mpox epidemics and advancing global public health initiatives.

## INTRODUCTION

The Mpox virus is emerging as a zoonotic disease of increasing concern due to its ability to infect humans. Primates have been identified as the primary reservoir of the virus; however, other mammals, including African rodents, have also been documented as potential carriers ^1^. Taxonomically, Mpox belongs to the family Poxviridae and the genus Orthopoxvirus (OPXV) ^2^. The molecular structure of Mpox is complex, with its virion consisting of a viral core, lateral bodies, an outer membrane, and a lipid envelope. Its genome comprises circular double-stranded DNA of approximately 197 kilobases (kb), encoding around 190 non-overlapping open reading frames (ORFs) ^3–5^. These ORFs encode various proteins that contribute to viral biology, including immunomodulation, host diversity, and pathogenicity.

The symptoms of Mpox infection vary, ranging from general symptoms such as fever and fatigue to characteristic skin lesions, and can lead to severe complications like pneumonia and encephalitis ^6^. The pathogenesis involves a complex interaction between the virus and the host, from viral entry to replication and dissemination throughout the body.

Mpox poses a significant challenge to global public health due to its capacity to cause epidemic outbreaks, with the most recent occurring in 2022 and 2024 ^7,8^. Therefore, developing effective vaccines is crucial for mitigating the impact of viral infection, and computational approaches for *in silico* peptide vaccine design have emerged as promising strategies ^9^. This study aims to identify and evaluate HTL epitopes in the Mpox virus to design a peptide vaccine capable of inducing specific immune responses that provide effective protection against the virus on a global scale. This work integrates molecular biology, bioinformatics, and computational modeling. The rapid identification and evaluation of potential vaccine candidates through computational approaches can accelerate vaccine development and improve preparedness for future Mpox outbreaks, making it a critical tool for preventing and controlling this emerging disease.

## RESULTS

### Surface protein selection

Following the annotation of the consensus genomes from the different Mpox clades, 17 proteins associated with the viral membrane and envelope were identified and selected for vaccine development. These proteins are depicted in the lower central part of Figure 1, with their coding genes highlighted in red. These proteins were chosen as key candidates for vaccine development and for use in generating potential vaccine epitopes.

**Figure 1.**
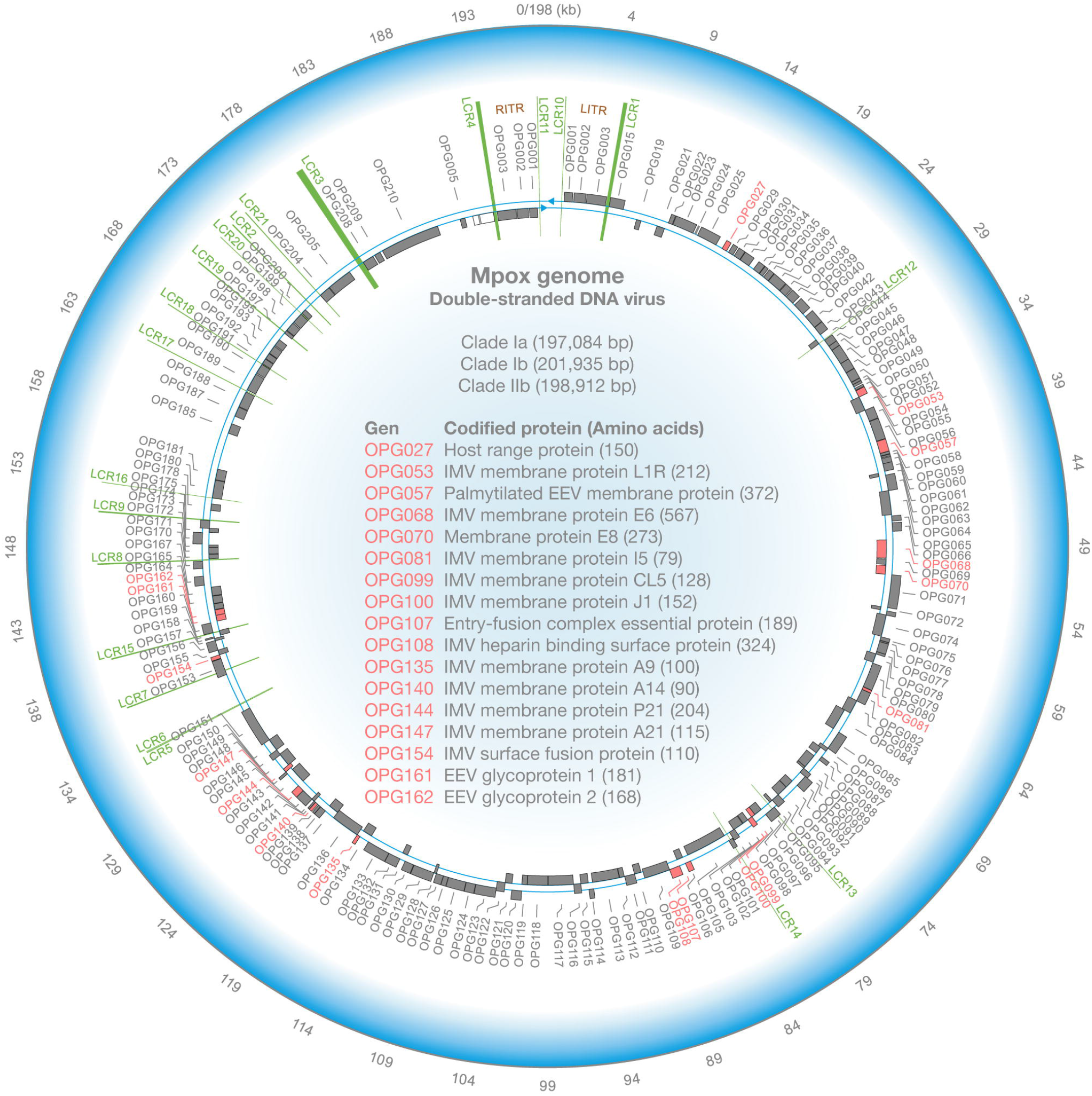
Flowchart illustrating the construction process of the Mpox vaccine.

### Selection of HLA-II sub-alleles

After retrieving HLA-II sub-allele frequencies from the Allele Frequency Database, sub-alleles with more than 5% prevalence in the global population were selected. The resulting alleles and their respective frequencies are presented in Supplementary Table 1, ensuring broad global and population coverage, with no populations excluded.

### Selection of vaccine epitopes by affinity

Using the IEDB MHCII Binding software, 50,040 potential epitopes were initially identified. These epitopes were evaluated with a length range of 12-18 amino acids. To ensure the quality and relevance of the selected epitopes, strict criteria were applied based on affinity score and binding rank with HLA-II. Only epitopes with an affinity score greater than 0.75 and a rank lower than 0.25 for each sub-allele were considered for this study.

As shown in Figure 2, the X-axis represents the epitopes, while the Y-axis displays the affinity scores obtained from the netMHCIIpan_EL 4.1 method. Each point is color-coded according to the specific sub-allele, allowing visualization of the variability in epitope interactions with different alleles. The size of each point indicates the rank, distinguishing strong from weak binders. Points with higher scores suggest a greater likelihood of epitope binding to the sub-allele, indicating higher potential efficacy in the immune response.

**Figure 2.**
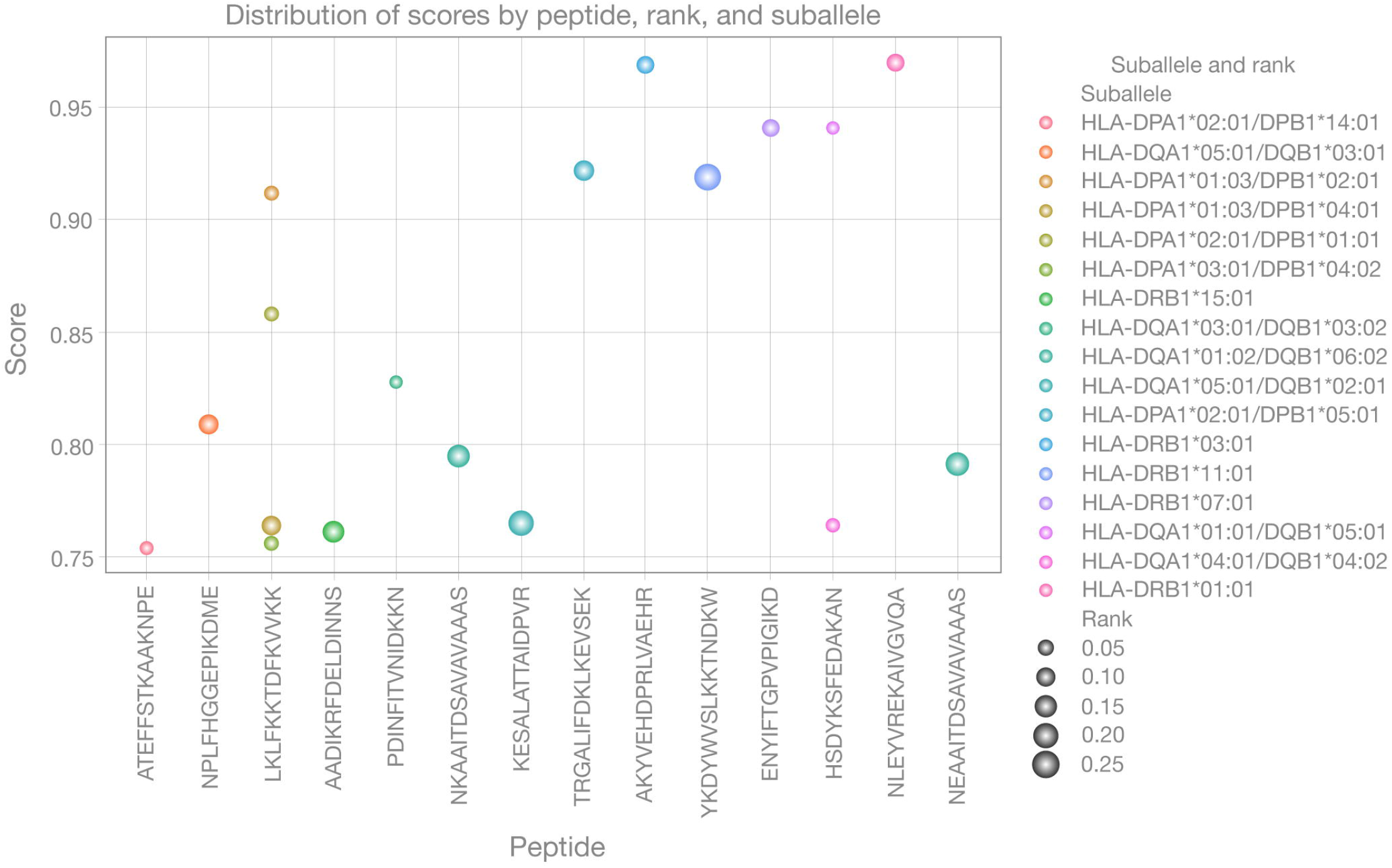
Functional annotation of the Mpox genome, a circular double-stranded DNA virus. The upper central section of the figure displays the consensus genome lengths for clades I, Ib, and IIb. The lower central section highlights the coding genes for the membrane and envelope proteins selected for this study, shown in red and labeled with their OPG (orthopoxvirus ortholog genes) nomenclature. Moving from the outer to the inner layers, the figure includes an inner ring representing the annotated genome, with genes of interest in red and non-interest genes in gray. Additionally, low-complexity regions (LCRs) are marked in green, along with terminal inverted repeat regions (TITRs) and internal terminal inverted regions (ITRs). The outer ring provides reference numbers representing the Mpox genome length, with a total reference of 198 kb.

### Evaluation of antigenicity, allergenicity, toxicity, stability, and homology

As shown in Supplementary Table 2, epitopes were filtered based on specific selection criteria: a) Antigenicity: The selected epitopes were evaluated using Vaxijen 2.0, with a threshold set at 0.4 for positive antigen predictions. A significant proportion of the selected epitopes exceeded this threshold, indicating high antigenic potential; b) Allergenicity: Using AllerTop v2.0, epitopes with allergenic potential were discarded. All selected epitopes were considered non-allergenic; c) Toxicity: Toxicity evaluation was conducted using ToxinPred, ensuring the absence of toxic properties. All selected epitopes were categorized as non-toxic; d) Stability: The stability of the epitopes was assessed using the ProtParam tool. Only epitopes with an instability index below 40.00 were retained, ensuring structural stability; and e) Homology: Homology with human proteins was evaluated using the NCBI BLAST tool, allowing the exclusion of homologous epitopes.

### Multi-epitope vaccine assembly

The vaccine assembly and its protein sequence are shown in Figure 3, where the 50S region of the *Mycobacterium* ribosomal protein L7/L12 was used as an adjuvant (red) and fused with the EAAAK linker (blue), followed by the selected epitopes (black) and their corresponding GPGPG linkers (turquoise). Finally, a histidine hexamer tag was added (orange).

**Figure 3.**
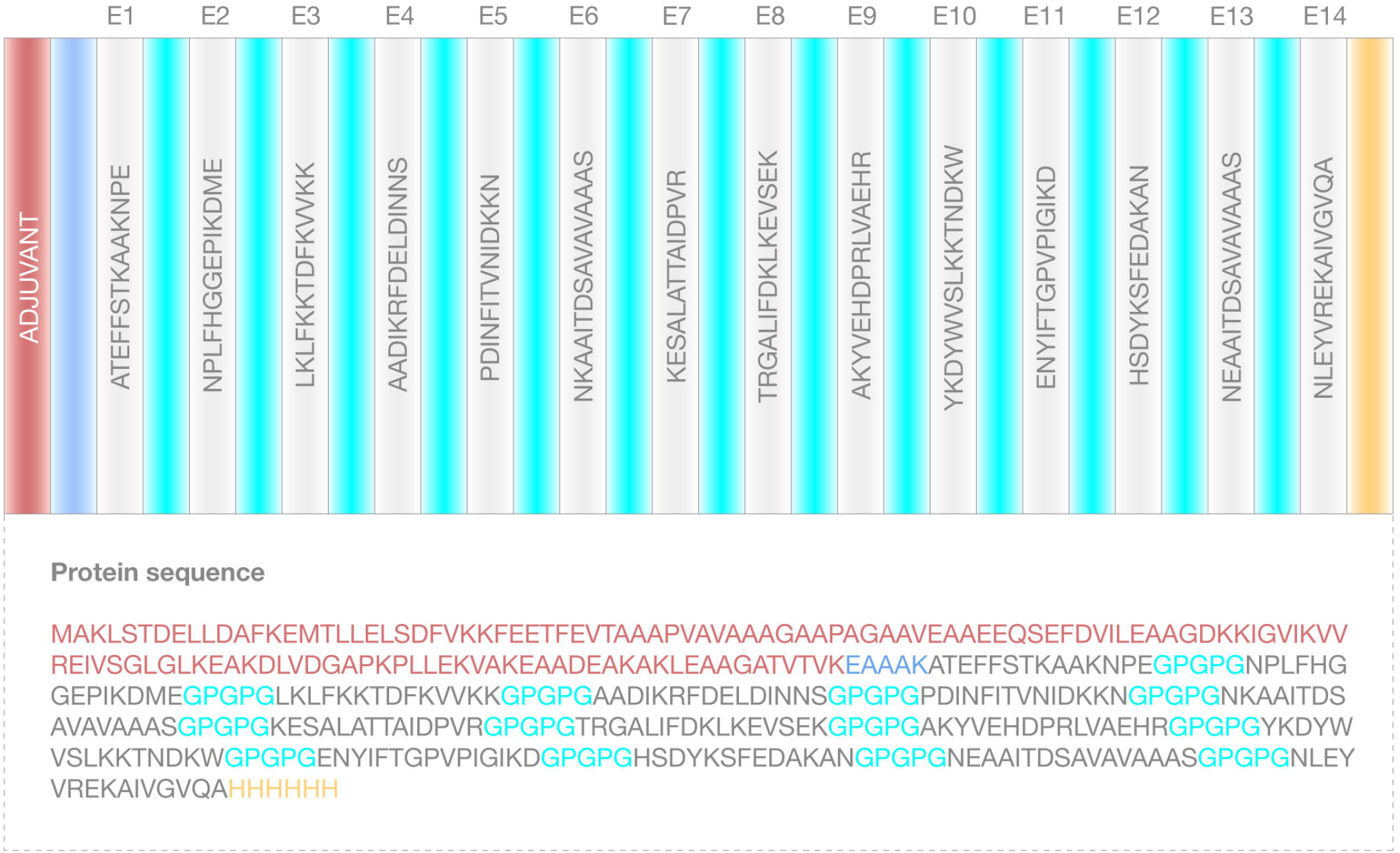
Dot plot showing the affinity of epitopes with suballeles according to the netMHCIIpan_EL 4.1 method, along with their corresponding “rank.”

### Immune System Simulation

The immune system simulation using C-ImmSim software provided detailed data on the immune response induced by the vaccine. In Figure 4A, cytokine and interleukin production is observed starting from day 0 of vaccination, indicating rapid activation of immune pathways. The central figure (D for “danger”) highlights the potential for a cytokine storm, a phenomenon that can trigger an uncontrolled inflammatory response. However, the results suggest a low likelihood of this event, reflecting proper immune regulation post-vaccination.

**Figure 4.**
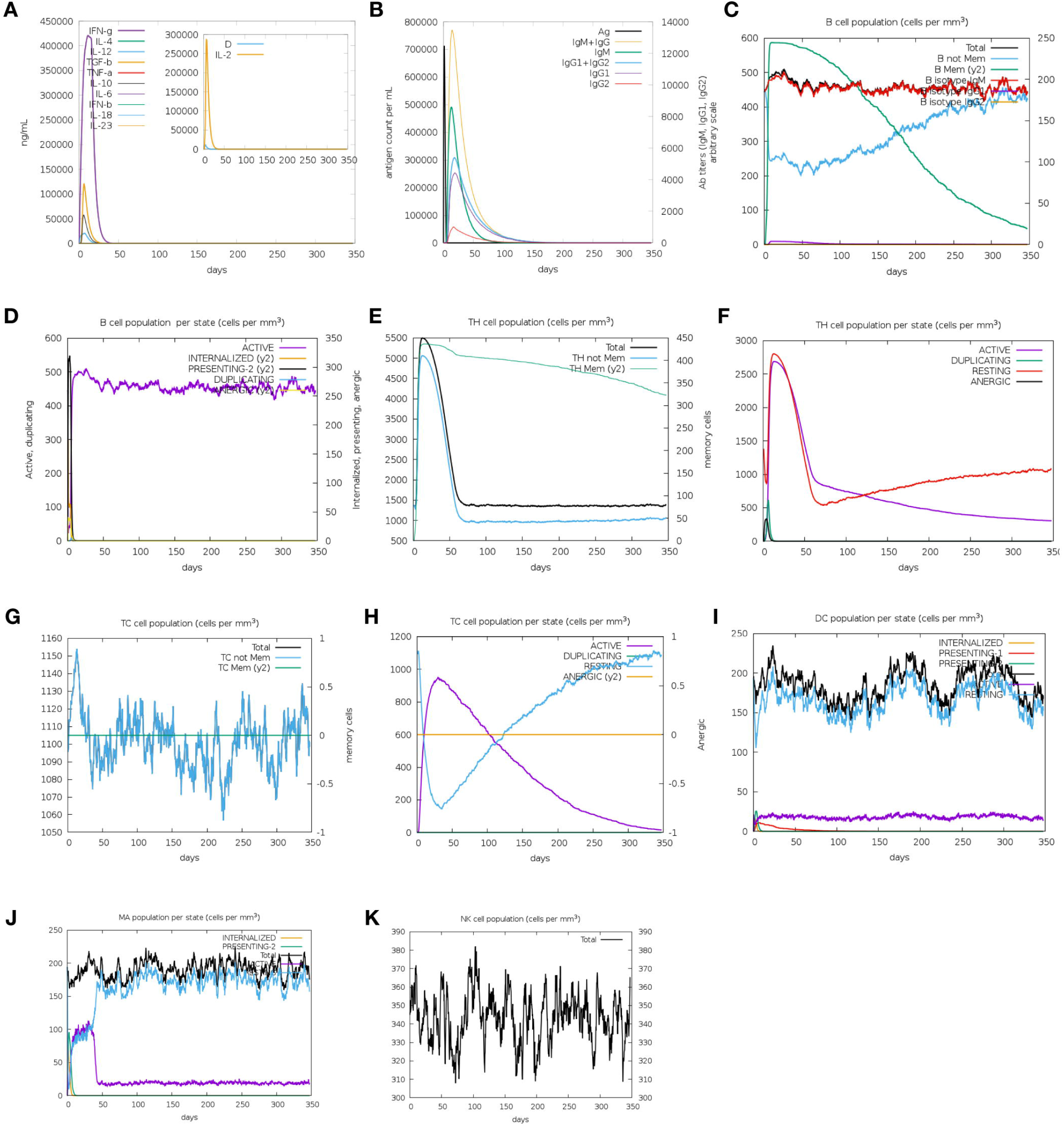
Assembly of the Mpox multi-epitope vaccine. Components include the adjuvant (red), EAAAK linker (blue), epitopes (black), GPGPG linkers specific for HTL cells (turquoise), and the histidine tag (HHHHHH) (orange).

In Figure 4B, the presence of the antigen (Ag) is detected from day 0, accompanied by increasing levels of immunoglobulins, with IgG dominating the secondary response, indicating effective and long-lasting immune memory. Figures 4C and 4D show the dynamics of memory B cells (y2) and non-memory B cells, which are activated from day 5, suggesting a robust and effective humoral response in the early stages of vaccine exposure.

Figures 4E and 4F depict the activation of HTLs, both memory and non-memory, starting on day 5 of exposure. Memory HTLs exhibit prolonged persistence, indicating a critical role in long-term adaptive immunity. Similarly, Figures 4G and 4H show the activation of cytotoxic T cells (Tc), following a pattern similar to that of HTLs but focused on the direct destruction of infected cells.

Additionally, simulations involving other key immune cells, such as dendritic cells (DCs), macrophage-activated killer cells (MA), and natural killer cells (NK), are illustrated in Figures 4I, 4J, and 4K. Dendritic cells, known for their ability to present antigens and activate both T and B cells, showed efficient activation following antigen exposure, highlighting their central role in initiating adaptive immunity. MA cells did not exhibit dysregulated responses, indicating well-controlled cytotoxic activity in this model. Lastly, NK cells, part of the innate immune response, demonstrated a rapid and effective response, contributing to the elimination of infected cells during the early phases of the immune response induced by the vaccine.

### 3D alignment and modeling

Figure 5A presents the predicted secondary structures of the vaccine, obtained using PSIPRED, while Figure 5B shows the three-dimensional structure of the multi-epitope vaccine modeled with I-TASSER. Initially, the structure had a GDT-HA of 0.55, RMSD (Root Mean Square Deviation) of -1.37, MolProbity of 3.146, Clash score of 6.2, Poor rotamers at 15.4%, and Ramachandran favored residues at 62.0%. After refinement, these values improved significantly: GDT-HA increased to 0.9163, RMSD decreased to 0.491, MolProbity dropped to 2.285, Poor rotamers drastically reduced to 1.0%, and the percentage of favored residues in the Ramachandran plot (Figure 5C) increased to 84.9%, indicating that most residues are in stable conformations. Additionally, 98.3% of residues were located in allowed regions, validating the structure’s quality. These improvements were visualized using PyMol v3.0.2, with epitopes and linkers highlighted in specific colors to facilitate interpretation and analysis.

**Figure 5.**
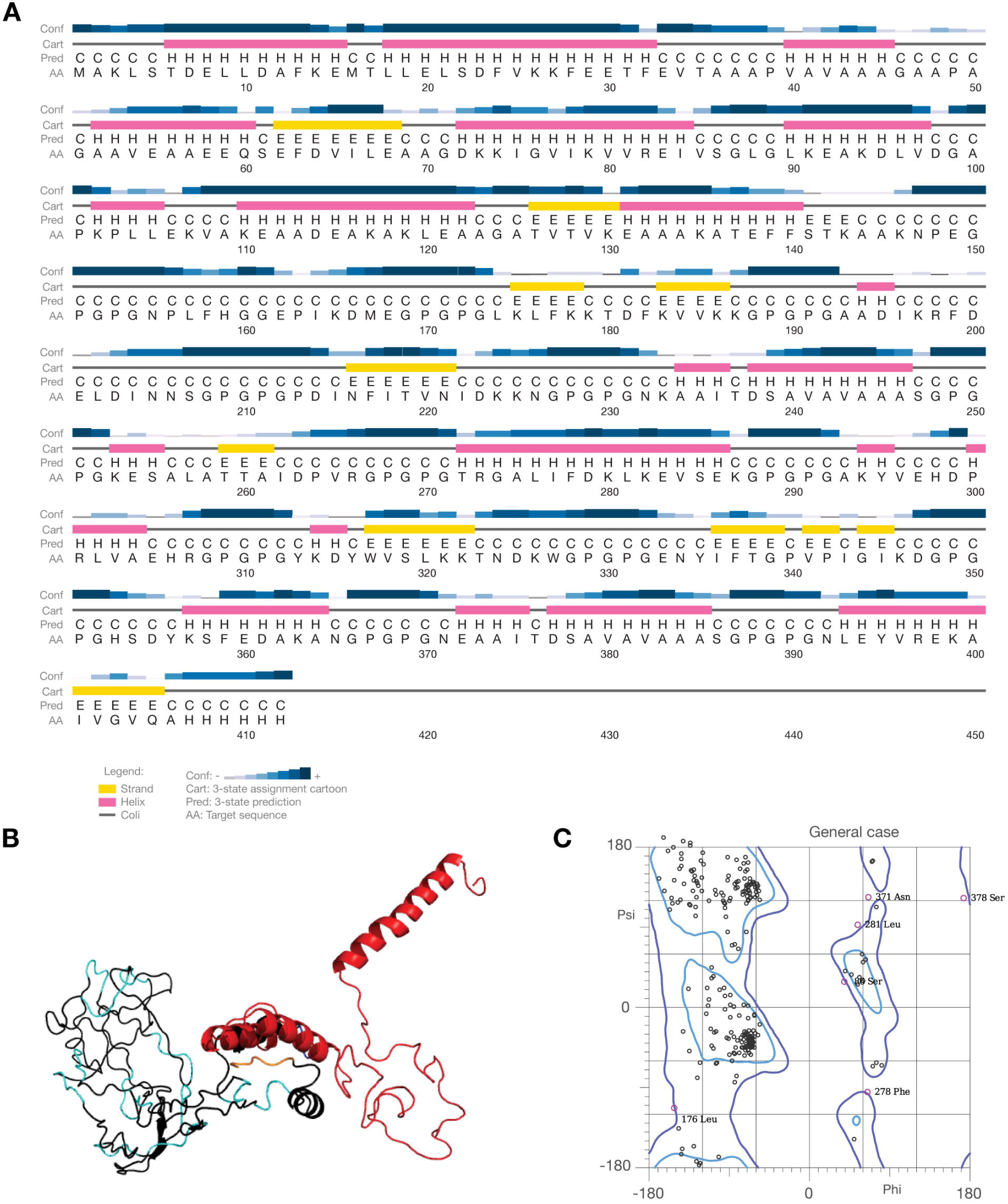
Simulation of the immune response to the Mpox vaccine, conducted using the C-ImmSim server.

### Population coverage analysis

The population coverage results showed that the vaccine achieved 99.8% global coverage. As shown in Figure 6, coverage was 98.9%, with an average hit (the average number of epitope-HLA matches recognized by the population) of 3.09, and a pc90 (the minimum number of epitope-HLA matches recognized by 90% of the population) of 2.03, indicating very high population coverage and efficient epitope recognition. Additionally, simulations by geographic regions demonstrated coverage ranging from 91.92% to 99.98%. These findings are detailed, with regional coverage represented in Figures 6A, 6B, and 6C.

**Figure 6.**
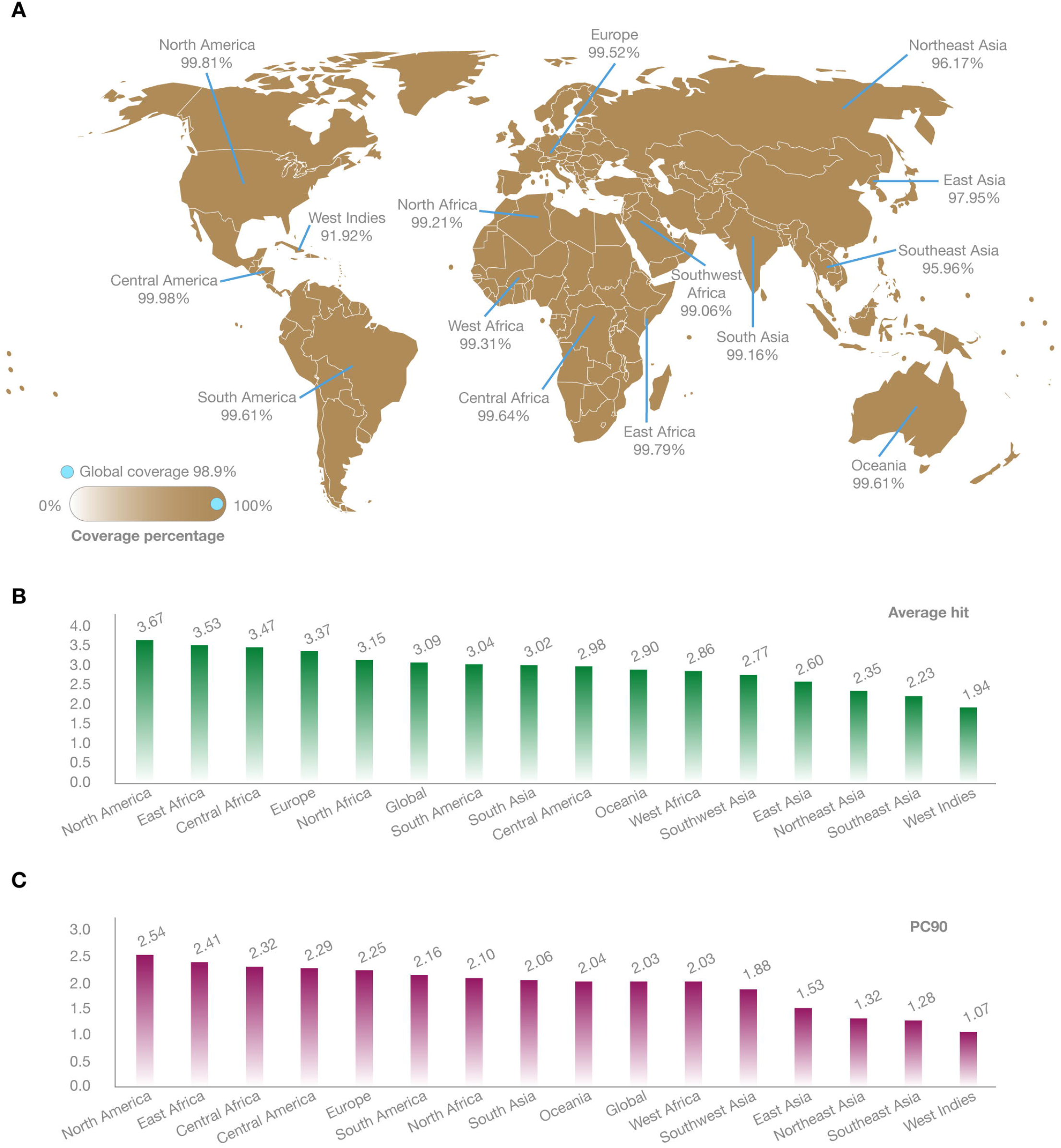
Subfigure A shows the PSIPRED evaluation for predicting the secondary structure of the Mpox vaccine. Subfigure B displays a 3D ribbon model representation of the protein structure, with the adjuvant in red, the EAAAK linker in blue, the epitopes in black, the GPGPG linkers specific for HTL cells in cyan, and the histidine tag (HHHHHH) in orange. Subfigure C presents the Ramachandran plot for protein validation.

### Prediction of vaccine interaction with immune system molecules

The HLA-DRB103:01, HLA-DPA103:01/DPB104:02, and HLA-DQA105:01/DQB103:01 sub-alleles were selected for homology modeling and subsequent molecular docking due to their high global prevalence. A total of 38 predicted structures were generated for each of the epitopes identified in the vaccine. Specifically, models where the ligand was positioned within the major groove were selected for free energy binding and Q-score calculations (Supplementary Table 3 and Figure 7). Figure 7 shows representative structures, highlighting the conformation between the two predominant alpha helices in the heterodimeric receptor with the analyzed ligands. All structures free energy binding values ranged from -1.51 to -46.26 kcal/mol, with several epitopes showing values more negative than -15 kcal/mol. Q-scores ranged from 0.012 to 0.11. Ligands E1, E4, E6, E10, and E13 exhibited the best poses and the highest Q-scores, ranging from 0.058 to 0.11.

**Figure 7.**
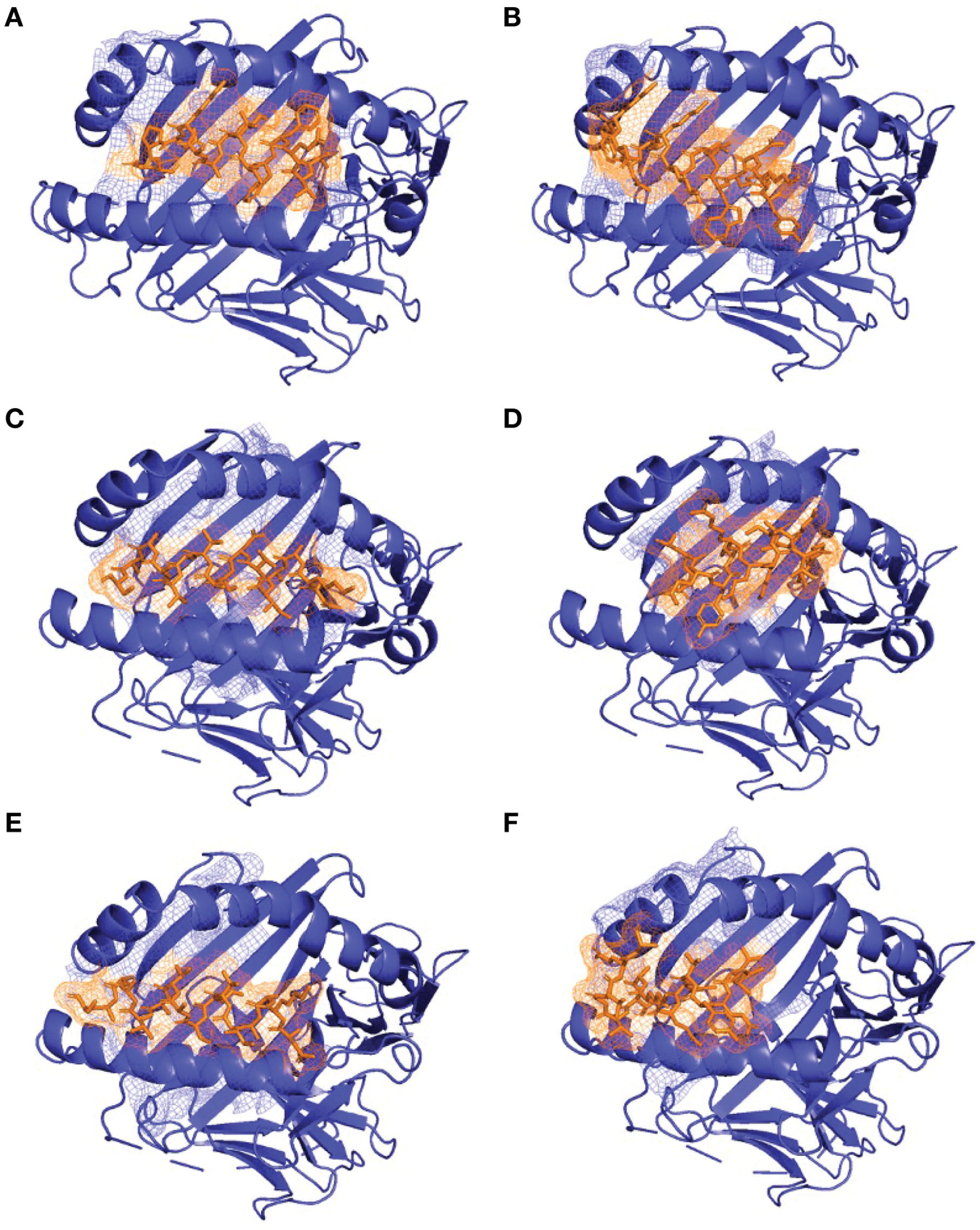
Global coverage prediction of the Mpox vaccine, generated using IEDB software.

## DISCUSSION

Today, there are several WHO-approved vaccines used against MPOX infection, including ACAM 2000 (Replicative Vaccine), LC16m8 (Semi-Replicative Vaccine) and MVA-BN (Non-Replicative Vaccine). These vaccines are based on attenuated or modified viruses and were designed to induce an immune response based on the replicative capacity of the attenuated virus into the host ^10^. However, computational resources can now be used to design a vaccine that exploits the potential of a more specific and adaptive multi-epitope response^11^. The *in silico* approach used in this study represents a significant innovation by designing a multiepitope vaccine with the capacity to maximize the immune response by targeting multiple viral regions. Unlike vaccines based on attenuated or modified viruses, this computational strategy allows the design of a vaccine with a robust immune response without the need to manipulate and inoculate live attenuated viruses, which can reduce risks for immunocompromised individuals^12^. Moreover, by targeting different areas of the virus, the proposed vaccine not only has the potential to protect the different clades of Mpox but also against other orthopoxviruses, which makes it a possible alternative to the current options, improving both immunological specificity and safety for the global population^13^.

The analysis performed with the NetMHCIIpan model allowed the identification of epitopes with high affinity for HLA-II molecules (Figure 2), important in the activation of the adaptive immune response^14^. This computational approach not only allows the selection of epitopes that induce robust immune responses but also provides a level of specificity that vaccines based on attenuated or modified viruses cannot achieve. The NetMHCIIpan model uses two key metrics: “rank” and “score.” The “rank” refers to the classification of the epitope relative to a reference set of sequences, used to categorize the strength of the epitope binding to HLA-II. According to established criteria, a rank below 0.5 is associated with strong binders (SB), while a rank below 2% is considered weak binders (WB) ^14^. In this study, a stricter selection threshold was applied, at <0.25. This ensures that the epitopes included in the analysis have the highest probability of acting as strong binders to HLA-II, which is crucial for vaccine design and other immunotherapeutic applications.

The “score” provides a direct measure of the likelihood that a peptide will be an HLA-II ligand and bind tightly to the complex. The score is expressed on a scale where values close to 1 indicate a high probability of effective binding ^14^. In this analysis, epitopes with a score greater than 0.75 were prioritized, suggesting that the ability to bind to HLA-II is possible while maintaining stable and durable interactions. This is particularly relevant when aiming to maximize the efficacy of an HTL-mediated immune response. By using score (>0.75) and rank (<0.25) values, the inclusion of highly immunogenic epitopes was guaranteed, enhancing the ability of the immune system to recognize different regions of the virus and generate a more targeted and specific response^14^.

A critical consideration in multi-epitope vaccine design is ensuring that the selected epitopes are not homologous to human proteins, which has important implications for the vaccine’s safety and efficacy ^15^. If the included epitopes share structural or functional similarity with endogenous proteins, the risk of the immune system failing to distinguish between viral antigens and self-proteins upon vaccine-mediated activation is high, potentially inducing the production of anti-homolog antibodies ^16^. This phenomenon could trigger autoimmune responses, leading to serious complications ^17^. To ensure high specificity, exhaustive homology tests were conducted during the vaccine assembly, ensuring that the homology coverage with human proteins did not exceed 40%. As shown in Supplementary Table 2, the proposed vaccine epitopes met this threshold, improving both the immunological safety profile and antiviral response efficacy ^15,18^. A lower coverage threshold could minimize the risk of autoimmunity but might also exclude important epitopes with moderate homology to human proteins that remain safe and effective. By using a 40% coverage, a balance between safety and efficacy is achieved, ensuring that the selected epitopes do not trigger a significant autoimmune response ^19,20^.

The success of a multi-epitope vaccine lies in ensuring that the selected epitopes have the appropriate length and structure for effective presentation by HLA-II molecules, which normally binds to peptides between 12 and 18 amino acids, so in this study epitopes with a length between 13 and 16 amino acids were selected (Figure 3). This length allows for an optimal fit in the HLA-II binding groove, facilitating a stable and effective interaction with the HLA-II complex ^21^. Additionally, this length ensures that the epitope can interact with multiple anchor sites within the HLA-II binding groove, enhancing the complex’s stability and promoting efficient antigen presentation ^22^. Epitopes of this length can also present the specific motifs and sequences necessary for recognition by T-cell receptors (TCR), optimizing HTL activation ^23^. This enhances the proposed vaccine interaction for inducing robust and specific immune responses, as it enables HTLs to coordinate and amplify adaptive immune responses, including B-cell activation and antibody production, highlighting the importance of long-lasting and effective immunity ^22,24^.

Moreover, when using the L7/L12 adjuvant derived from *Mycobacterium*—especially from BCG or Bacillus Calmette-Guérin—we emphasize its role as a potent enhancer of T-cell activation through B-cells, capitalizing on its robust immunostimulatory properties observed in interactions with DCs, as evidenced by antigen recognition in Figure 4A ^25^.

When this proposed vaccine is phagocytosed, it induces marked maturation of these cells, characterized by the high expression of co-stimulatory molecules like CD40, B7.1, and B7.2, which are necessary to facilitate superior antigen presentation and subsequent T-cell activation ^26,27^. Moreover, coculture studies reveal that DCs exposed to BCG induce significant upregulation of activation markers like CD25, CD54, and CD71 on CD4+ and CD8+ T cells, accompanied by intensified secretion of IL-2, IL-10, and IFN-γ, as seen in Figure 4C. Mechanistically, blocking studies targeting B7.1, B7.2, or IL-12 strongly attenuate IFN-γ production, further validating the critical role of these pathways in T-cell activation. Thus, the L7/L12 adjuvant stands out as a promising candidate for enhancing B-cell-mediated T-cell activation, potentially advancing its application in Mpox vaccine development ^26–28^.

The GPGPG linkers used in the proposed vaccine are specific for HTLs, playing a crucial role in the immune response. These linkers are strategically designed to optimize antigen presentation to HTLs via HLA-II molecules. The repetitive structure of GPGPG provides flexibility and stability to the vaccine, enabling specific and efficient interaction with HTL receptors ^27,29,30^. This interaction facilitates HTL activation, which in turn promotes coordinated immune responses, including cytokine production and activation of B and cytotoxic cells. Furthermore, GPGPG linkers not only improve antigen presentation but can also modulate the duration and intensity of the adaptive immune response, ensuring robust and sustained protection against specific pathogens. On the other hand, the histidine tag (HHHHHH) used in the assembly (Figure 3) plays a crucial role in vaccine design due to its unique properties that enhance the formulation’s efficacy. The utility of this histidine sequence lies in its function as an affinity tag, facilitating the purification of the recombinant antigen during production, ensuring high purity and concentration of the final antigen ^31^.

Immunological simulations offer a crucial tool for understanding how the body responds to infections or vaccines, allowing us to observe complex dynamics without resorting to invasive testing. In our study, Figure 4A shows how cytokines and interleukins respond after exposure, staying within normal levels to avoid an uncontrolled immune response ^32^. This is critical because, as observed, the danger rate (D) is positioned below the levels of IL-2, a key cytokine involved in T cell activation and immune regulation ^33,34^. Controlling this molecule is essential since overproduction of cytokines can lead to a cytokine storm, triggering systemic inflammation and tissue damage, thus compromising the safety of the vaccinated individual ^35^.

In Figure 4B, the dynamics of IgM and IgG production in response to the antigen are shown, which is key for humoral defense. The antigen is recognized around day 5, after which immunoglobulin production begins. Early IgM production provides rapid initial defense, while the later appearance of IgG ensures more specific and long-lasting protection ^36,37^. This behavior is important because a quick and efficient response not only ensures pathogen neutralization but also aids in the development of immune memory, as early antigen recognition followed by the production of effective antibodies guarantees a robust response against future exposures ^38,39^.

Figures 4C, 4E, and 4G demonstrate the production of memory cells, specifically memory B lymphocytes, memory TH cells, and memory TC cells. These cells play a vital role in long-term immunity. The persistence of these memory cell populations is essential as they enable the immune system to respond quickly and efficiently if the organism is re-exposed to the same pathogen ^40,41^. Memory TC cells show stability over time, indicating a durable immune response. This stability is crucial to ensure that in future exposures, the immune system can eliminate infected cells more rapidly, thereby limiting the severity of the infection ^41^.

Finally, Figures 4D, 4F, and 4H detail the activation and functionality states of lymphocytes, revealing important phases such as activation, duplication, and anergy. These processes are critical to ensure that the immune system responds in a controlled manner ^41^. In particular, the internalized B lymphocytes presenting antigens in Figure 4D stand out for their essential role in activating other immune cells, a key step in orchestrating a complete adaptive immune response ^42,43^. Similarly, dendritic cells (Figure 4I), macrophages (Figure 4J), and NK cells (Figure 4K) emphasize the importance of cellular cooperation in the immune response, acting as mediators in both innate and adaptive immunity ^43^.

The results obtained demonstrate a significant improvement in the structural quality of the multi-epitope vaccine model. The increase in GDT-HA to 0.9163 reflects greater accuracy in the global superposition of the model concerning the reference structure, while the reduction in RMSD to 0.491 indicates less average atomic deviation, suggesting a conformation closer to the native state. The decrease in MolProbity to 2.285 and the percentage of poor rotamers to 1.0% reinforces structural stability, showing a reduction in stereochemical errors in the side-chain conformations of the residues ^44,45^. Although the Clash score remained at 13.0, which could suggest slightly unfavorable interactions between nearby atoms, the notable improvement in the percentage of favored residues in the Ramachandran plot, as seen in Figure 5C (which increased to 84.9%), along with 98.3% of residues located in allowed regions, confirms that most residues adopt energetically favorable conformations ^44,46^. These results consolidate the robustness and feasibility of the proposed vaccine model.

The design of vaccines that are effective for most of the global population represents a significant challenge, as genetic diversity and differences in HLA alleles across various geographical regions can influence the immune response. To develop vaccines that generate an effective immune response on a global scale, it is essential to assess the population coverage of the proposed epitopes ^47^. To ensure this coverage, the proposed vaccine achieved a remarkable result in global population coverage, with a success rate of 98.9%, as shown in Figure 6. Furthermore, it is revealed that the average number of epitopes recognized by the population is 3.09 (Average hit), indicating that at least 98.9% of the population will recognize an average of 3 epitopes from the proposed vaccine ^48^. Additionally, the vaccine presented a PC90 value of 2.03, meaning that at least 90% of HLA alleles globally will recognize at least 2 epitopes from the proposed vaccine ^48^. These results demonstrate that the epitopes proposed for the design of this vaccine will be recognized by different HLA alleles, ensuring the vaccine’s antigenicity. When evaluating various population points, as observed in Figure 6, Central America has the highest combined exposure to MHC-I and MHC-II alleles, reaching 99.98%, while Northern Asia shows the lowest population coverage at 96.17% ^48,49^.

To support and deepen the analysis initiated with NetMHCIIpan, a molecular docking analysis was performed with each of the epitopes previously identified and incorporated into the vaccine design. The predicted poses for nearly all epitopes showed the peptides positioned near or within the peptide-binding groove of the modeled HLA sub alleles receptors. The flexible groove in the human MHC-II structure can accommodate 13 to 18 amino acids, which is necessary for the correct positioning of the structure within this domain, ensuring stable binding and effective presentation to T cells ^50^. The predicted general binding energy values indicate stable conformations between receptors and ligands, with more negative energies observed for ligands positioned across the groove and higher Q-scores compared to those with more distinguishable deviations from the template structure. Overall, all the epitopes presented negative binding energies, suggesting favorable conditions for forming complexes with MHC-II and successfully presenting different antigens. This approach differs from similar studies, which report energy values around -7 to -12 kcal/mol, in some cases using global energies as an indicator of affinity between the MHC-II and the complete vaccine proposed ^51,52^. Despite the variation in free energy binding with the three suballeles used as receptors, it is important to note that HLA class II molecules exhibit varying preferences for peptides due to polymorphisms within the binding groove. These differences are critical in determining immune responses, disease susceptibility, and vaccine efficacy ^53^. Therefore, other prevalent HLA receptors may possess differential adaptive capabilities for these epitopes, potentially improving overall antigen presentation and triggering a more effective immune response.

## LIMITATIONS AND CONCLUSIONS

The primary limitation of this study is the absence of *in vitro* and *in vivo* validation for the results obtained through computational tools. While the *in silico* approach enables the prediction of epitopes with high immunogenic potential, these findings must be confirmed through laboratory experiments. *In vitro* assays, such as tests for epitope binding to HLA-II molecules and T-cell activation, are crucial to verify the actual ability of the selected epitopes to induce an immune response. Similarly, *in vivo* studies in animal models are essential to evaluate the safety, immunogenicity, and efficacy of the vaccine in a complex biological system. Without these validations, predicting how the multi-epitope vaccine will perform in living organisms is challenging, and any assumptions about its effectiveness in humans remain speculative.

Nevertheless, this study represents a significant advancement in vaccine engineering, using an *in silico* approach to design a multi-epitope vaccine targeting the surface proteins of the Mpox virus. Through the application of advanced bioinformatics tools, candidate epitopes were rigorously identified, selected, and evaluated, demonstrating high affinity for HLA-II complexes and ensuring proper presentation to HTLs. These epitopes were chosen not only for their positive antigenicity but also for their lack of allergenicity and toxic potential.

Computational simulations showed that the designed vaccine can induce a robust and memory-based immune response, which is crucial for long-term efficacy. The 3D modeling confirmed the stable structure of the vaccine and the optimal exposure of the epitopes to HTLs, facilitating efficient interaction and a coordinated immune response.

This study underscores the transformative potential of bioinformatics in the development of effective and personalized vaccines, highlighting its strategic importance in global efforts to prepare for and respond to emerging diseases.

## METHODS

To develop an effective multi-peptide-based vaccine against Mpox, various bioinformatics techniques were implemented using advanced computational methods, as shown in the flowchart in Figure 8.

**Figure 8.**
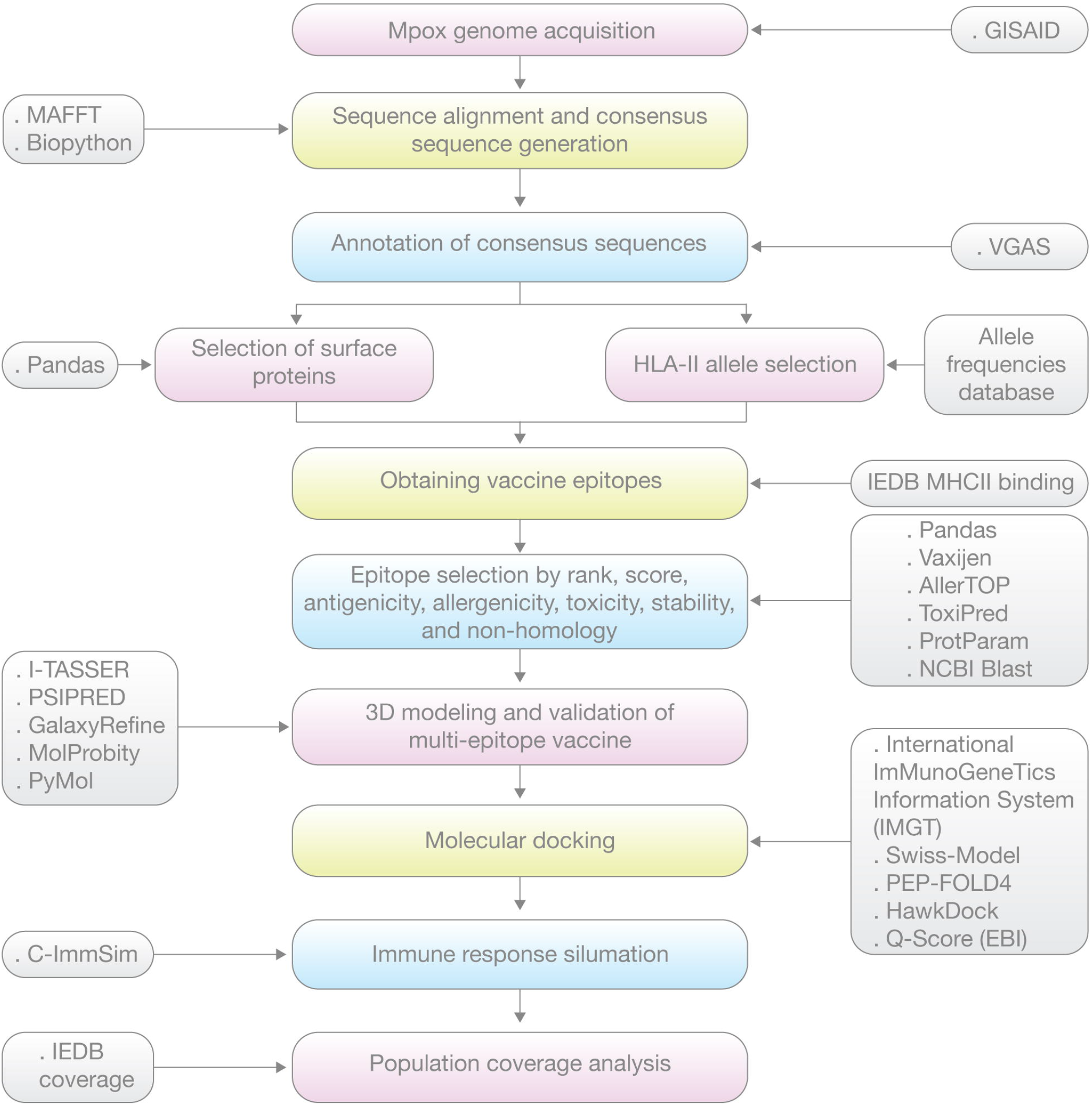
Representative docking poses of the HLA receptors with specific vaccine epitopes. The predicted models show the HLA receptors (purple) in complex with different epitopes/ligands (orange). A) HLA-DRB103:01 - E1. B) HLA-DRB103:01 - E10. C) HLA-DPA103:01/DPB104:02 - E6. D) HLA-DPA103:01/DPB104:02 - E13. E) HLA-DQA105:01/DQB103:01 - E6. F) HLA-DQA105:01/DQB103:01-E13.

### Sequence acquisition

A total of 154 Mpox virus sequences were retrieved from the GISAID EpiPoxTM database (https://gisaid.org/), representing clades Ia, Ib, and IIb. Filtering criteria were applied to ensure that only complete genomes (>196,000 bp) were included ^54^.

To maintain the genetic diversity of the samples, sequences from various geographic regions and viral clades were selected and analyzed using the Nextclade v3.8.2 tool (https://clades.nextstrain.org/) ^55^. As detailed in Supplementary Table 4 below, the geographic distribution of the samples is presented alongside their respective clades and lineages.

### Sequence alignment and consensus sequence generation

After collecting the 154 sequences, a multiple sequence alignment was performed using the MAFFT software for each clade (https://mafft.cbrc.jp/alignment/server/index.html), resulting in three alignment files in multi-fasta format (clades Ia, Ib, IIb) ^56^. The consensus sequence was generated using the Biopython V1.84 library (https://biopython.org/) and its AlignIO module ^57^. This process involved reading the alignment file and determining the consensus sequence by identifying the most frequent nucleotide base at each position across all aligned sequences.

### Functional annotation of the consensus sequence

The functional annotation of the consensus sequences was performed using the VGAS tool (http://guolab.whu.edu.cn/vgas/). This tool provided the coordinates of predicted genes, the nucleotide sequences of each potential gene, and their respective proteins ^58^. Proteins located on the viral surface were selected using the Pandas library (https://pandas.pydata.org/), considering their relevance in host-pathogen interactions and the development of antiviral therapies ^59^.

### HLA-II allele selection

HLA-II alleles with a global prevalence of more than 5% were selected, specifically from the HLA-DR, HLA-DQ, and HLA-DP classes. Allele frequencies were obtained from the Allele Frequencies Database (https://www.allelefrequencies.net/default.asp). The frequencies of alleles were collected by country, and the most prevalent alleles globally were calculated and selected ^60^.

### Epitope identification and analysis

MHC-II binding predictions from the Immune Epitope Database and Analysis Resource (IEDB) (https://www.iedb.org/) were used for epitope identification. Previously annotated viral surface proteins were analyzed with a peptide length range of 12-18 amino acids, alongside the previously selected HLA-II alleles ^61^.

### Vaccine epitope selection

After identifying potential vaccine epitopes, the Pandas library was used to select those with an affinity score greater than 0.75 and a rank lower than 0.25, using the netMHCIIpan_EL 4.1 method from IEDB. This method predicts the binding affinity of epitopes to HLA-II sub-alleles based on their interaction probabilities. The selected affinity values indicate a high likelihood of stable interaction between the epitope and the allele, suggesting strong binding, which is crucial for effectively activating an immune response ^59^.

Subsequently, filters for antigenicity, allergenicity, toxicity, stability, and homology with human proteins were applied. To evaluate the antigenicity of the epitopes, the Vaxijen 2.0 tool (https://www.ddg-pharmfac.net/vaxijen/VaxiJen/VaxiJen.html) was used with a default threshold of 0.4 to ensure a positive prediction of antigenicity, which is essential for stimulating the immune system ^62^. Allergenicity was assessed using the AllerTop v2.0 tool (https://www.ddg-pharmfac.net/AllerTOP/method.html), and epitopes with allergenic potential were excluded ^63^. Additionally, a toxicological analysis was performed using the ToxinPred tool (https://webs.iiitd.edu.in/raghava/toxinpred/algo.php) to identify and eliminate potentially toxic peptides ^64^. The stability of the peptides was then evaluated using the ProtParam Tool (https://web.expasy.org/protparam/), which predicts the instability index of the peptides. An instability index greater than 40.00 indicates that a peptide is unstable, while an index below 40.00 suggests stability. These criteria ensure the selection of epitopes with optimal properties for designing a vaccine capable of activating HTLs ^65^. Finally, the homology of the epitopes with human proteins was assessed using the NCBI BLAST server (https://blast.ncbi.nlm.nih.gov/Blast.cgi) to exclude those with significant similarities that could compromise the vaccine’s safety and specificity. Epitopes with homology coverage below 40% were selected to ensure they do not share significant similarities with human proteins ^66^.

### Vaccine assembly

To enhance the immunogenicity of the selected epitopes, a specific adjuvant was used. In this study, the 50S region of the ribosomal protein L7/L12 from the *Mycobacterium* genus was employed, obtained from the NCBI database (https://www.ncbi.nlm.nih.gov/protein) under accession number WP_003403353.1. This region was fused with an EAAAK linker to provide flexibility and structural stability, followed by the selected epitopes and specific linkers for HTL cell epitopes, consisting of the GPGPG sequence. Finally, a hexameric histidine tag was added to facilitate purification and further analysis of the chimeric protein ^27^.

### 3D modeling of the multi-epitope vaccine

The I-TASSER tool (https://zhanggroup.org/I-TASSER/) was used to model the protein structure, predicting the protein’s structure based on its amino acid sequence using an iterative template-based assembly approach ^67^. Subsequently, PSIPRED (http://bioinf.cs.ucl.ac.uk/psipred/) was used to generate a visual representation of the predicted secondary structure, showing α-helices, β-sheets, and coil regions, enabling an interpretation of protein conformation ^68^. Next, GalaxyRefine (https://galaxy.seoklab.org/) was used to improve the accuracy and quality of the modeled structure ^46^. The refined model was evaluated using a Ramachandran plot through MolProbity (http://molprobity.biochem.duke.edu/), assessing the quality of polypeptide chain conformations based on the φ (phi) and ψ (psi) torsion angles of peptide bonds ^69^. Finally, the resulting PDB file was visualized using PyMol v3.0.2 (https://pymol.org/) ^70^.

### Immune response simulation

The C-ImmSim web server (https://kraken.iac.rm.cnr.it/C-IMMSIM/index.php) was used for immune response simulation. This server predicts, based on the vaccine sequence, the development of memory and non-memory B cells, the time required for the vaccine to induce immunogenicity, the behavior of immunoglobulins and pro-inflammatory factors, and the vaccine’s effectiveness in generating an immune response ^71^. C-ImmSim employs advanced computational models to simulate the adaptive immune response, providing a comprehensive evaluation of the designed vaccine’s immunogenic potential ^71^.

### Population coverage analysis

Population coverage for the specific vaccine was analyzed using the dedicated tool provided by the IEDB web server (https://www.iedb.org/) ^49^. Several simulations were performed using MHC-I and MHC-II epitope data to calculate results such as projected population coverage, average hit (the average number of epitope/HLA combinations recognized by the population), and pc90 (the minimum number of epitope/HLA combinations recognized by 90% of the population) ^49^. These simulations were conducted globally and by geographic regions, which were divided into 15 areas: North America, Central America, South America, Northern Africa, Central Africa, Western Africa, Eastern Africa, Europe, Oceania, Northeast Asia, South Asia, Southeast Asia, East Asia, Southwest Asia, and the West Indies ^49^.

### Molecular docking

For the docking predictions, the sequences of HLA sub-alleles (acting as receptor molecules) were obtained from the International ImMunoGeneTics Information System (https://www.ebi.ac.uk/ipd/imgt/hla/), and their structures were modeled using the Swiss-Model server (https://swissmodel.expasy.org/). The epitopes forming the vaccine structure, used as ligands, were modeled using the PEP-FOLD4 server with default settings, except for the ionic strength, which was adjusted to 150 mM ^72,73^. The predictions of binding poses and energies between the HLA complexes and the vaccine epitopes were carried out using the HawkDock server ^74,75^. This server integrates Molecular Mechanics/Generalized Born Surface Area (MM/GBSA) in a second step to calculate the free energy of selected structures, providing an initial estimate of binding affinities and complex stability. The final models were selected based on their binding poses, energies, and Q-Scores (https://www.ebi.ac.uk/msd-srv/ssm/) ^76^, compared to the reference structure. The Q-score, which ranges from 0 to 1, is used to evaluate the superposition of two proteins at the secondary structure level and was selected as a key indicator for determining the optimal spatial prediction of the peptide within the HLA domains.

## Acknowledgments

This work was supported by Universidad Internacional SEK (UISEK) and Universidad de Las Américas (UDLA) from Quito, Ecuador. Additionally, we would like to thank María Fernanda Gutiérrez, PhD for her suggestions, which contributed to improving the clarity and accuracy of the final version of this manuscript.

## Author Contribution

A.H.Y. and S.R.O. conceived the subject and the conceptualization of the study. S.R.O. and J.R.R.I. did data curation and supplementary data. S.R.O., J.R.R.I., J.D.A.E., J.E.E., J.C.N., A.L.C., and A.H.Y. gave conceptual advice and valuable scientific input. A.H.Y., J.D.A.E., and J.E.E. supervised the project. A.L.C. did a funding acquisition. Finally, all authors reviewed and approved the manuscript

## Data availability

All data generated or analyzed during this study are included in this published article (and its Supplementary Information files).

## Bibliography

1. Doty, J. B. et al. Assessing Monkeypox Virus Prevalence in Small Mammals at the Human-Animal Interface in the Democratic Republic of the Congo. Viruses 9, (2017).

2. Alakunle, E. et al. A comprehensive review of monkeypox virus and mpox characteristics. Front. Cell. Infect. Microbiol. 14, 1360586 (2024).

3. Huang, Y., Mu, L. & Wang, W. Monkeypox: epidemiology, pathogenesis, treatment and prevention. Signal Transduct. Target. Ther. 7, 373 (2022).

4. Luna, N. et al. Phylogenomic analysis of the monkeypox virus (MPXV) 2022 outbreak: Emergence of a novel viral lineage? Travel Med. Infect. Dis. 49, 102402 (2022).

5. Viruela símica, un reto para la salud pública mundial. http://scielo.sld.cu/scielo.php?pid=S1684-18242022000400637&script=sci_arttext.

6. Sun, Y., Nie, W., Tian, D. & Ye, Q. Human monkeypox virus: Epidemiologic review and research progress in diagnosis and treatment. J. Clin. Virol. 171, 105662 (2024).

7. Kröger, S. T. et al. Mpox outbreak 2022: an overview of all cases reported to the Cologne Health Department. Infection 51, 1369–1381 (2023).

8. Cevik, M. et al. The 2023 - 2024 multi-source mpox outbreaks of Clade I MPXV in sub-Saharan Africa: Alarm bell for Africa and the World. Int. J. Infect. Dis. 146, 107159 (2024).

9. Kanampalliwar, A. M. Reverse vaccinology and its applications. Methods Mol. Biol. 2131, 1–16 (2020).

10. Reina, J. & Iglesias, C. Vaccines against monkeypox. Med Clin (Barc) 160, 305–309 (2023).

11. Grabenstein, J. D. & Hacker, A. Vaccines against mpox: MVA-BN and LC16m8. Expert Rev. Vaccines 23, 796–811 (2024).

12. Morino, E. et al. Mpox neutralizing antibody response to lc16m8 vaccine in healthy adults. NEJM evid. 3, EVIDoa2300290 (2024).

13. Sadeghi, Z., Fasihi-Ramandi, M. & Bouzari, S. Evaluation of immunogenicity of novel multi-epitope subunit vaccines in combination with poly I:C against Brucella melitensis and Brucella abortus infection. Int. Immunopharmacol. 75, 105829 (2019).

14. Reynisson, B., Alvarez, B., Paul, S., Peters, B. & Nielsen, M. NetMHCpan-4.1 and NetMHCIIpan-4.0: improved predictions of MHC antigen presentation by concurrent motif deconvolution and integration of MS MHC eluted ligand data. Nucleic Acids Res. 48, W449–W454 (2020).

15. Salahlou, R. et al. Development of a novel multi-epitope vaccine against the pathogenic human polyomavirus V6/7 using reverse vaccinology. BMC Infect. Dis. 24, 177 (2024).

16. Haltaufderhyde, K. et al. Immunoinformatic risk assessment of host cell proteins during process development for biologic therapeutics. AAPS J. 25, 87 (2023).

17. Greinacher, A. et al. Insights in ChAdOx1 nCoV-19 vaccine-induced immune thrombotic thrombocytopenia. Blood 138, 2256–2268 (2021).

18. Shawan, M. M. A. K. et al. Advances in Computational and Bioinformatics Tools and Databases for Designing and Developing a Multi-Epitope-Based Peptide Vaccine. Int. J. Pept. Res. Ther. 29, 60 (2023).

19. Ahmed, M. H. et al. An immuno-informatics approach for annotation of hypothetical proteins and multi-epitope vaccine designed against the Mpox virus. J. Biomol. Struct. Dyn. 42, 5288–5307 (2024).

20. Toussaint, N. C. & Kohlbacher, O. Towards in silico design of epitope-based vaccines. Expert Opin. Drug Discov. 4, 1047–1060 (2009).

21. Castelli, F. A. et al. HLA-DP4, the most frequent HLA II molecule, defines a new supertype of peptide-binding specificity. J. Immunol. 169, 6928–6934 (2002).

22. Josephs, T. M., Grant, E. J. & Gras, S. Molecular challenges imposed by MHC-I restricted long epitopes on T cell immunity. Biol. Chem. 398, 1027–1036 (2017).

23. Gras, S. et al. Reversed T cell receptor docking on a major histocompatibility class I complex limits involvement in the immune response. Immunity 45, 749–760 (2016).

24. Nielsen, M., Lundegaard, C. & Lund, O. Prediction of MHC class II binding affinity using SMM-• align, a novel stabilization matrix alignment method. BMC Bioinformatics 8, 238 (2007).

25. Lee, S. J. et al. A potential protein adjuvant derived from Mycobacterium tuberculosis Rv0652 enhances dendritic cells-based tumor immunotherapy. PLoS ONE 9, e104351 (2014).

26. Liu, T., Shi, K. & Li, W. Deep learning methods improve linear B-cell epitope prediction. BioData Min. 13, 1 (2020).

27. Zakaria, M. N. Z., Aththar, A. F., Fai, M. & Hamami, S. M. A. Archive of SID. ir. (2024).

28. Sicard, T., Kassardjian, A. & Julien, J.-P. B cell targeting by molecular adjuvants for enhanced immunogenicity. Expert Rev. Vaccines 19, 1023–1039 (2020).

29. Behbahani, M., Moradi, M. & Mohabatkar, H. In silico design of a multi-epitope peptide construct as a potential vaccine candidate for Influenza A based on neuraminidase protein. In Silico Pharmacol 9, 36 (2021).

30. Ullah, A. et al. Bioinformatics and immunoinformatics approach to develop potent multi-peptide vaccine for coxsackievirus B3 capable of eliciting cellular and humoral immune response. Int. J. Biol. Macromol. 239, 124320 (2023).

31. Singh, M. et al. Effect of N-terminal poly histidine-tag on immunogenicity of Streptococcus pneumoniae surface protein SP0845. Int. J. Biol. Macromol. 163, 1240–1248 (2020).

32. Bülow Anderberg, S. et al. Increased levels of plasma cytokines and correlations to organ failure and 30-day mortality in critically ill Covid-19 patients. Cytokine 138, 155389 (2021).

33. Castiglione, F. & Bernaschi, M. C-immsim: playing with the immune response. … on mathematical theory of networks and … (2004).

34. Śledzińska, A. et al. Regulatory T Cells Restrain Interleukin-2- and Blimp-1-Dependent Acquisition of Cytotoxic Function by CD4+ T Cells. Immunity 52, 151-166.e6 (2020).

35. Jiang, Y. et al. Cytokine storm in COVID-19: from viral infection to immune responses, diagnosis and therapy. Int. J. Biol. Sci. 18, 459–472 (2022).

36. Perelson, A. S., Goldstein, B. & Rocklin, S. Optimal strategies in immunology III. The IgM-IgG switch. J. Math. Biol. 10, 209–256 (1980).

37. Rogier, E. et al. Antibody dynamics in children with first or repeat Plasmodium falciparum infections. Front Med (Lausanne) 9, 869028 (2022).

38. Isho, B. et al. Persistence of serum and saliva antibody responses to SARS-CoV-2 spike antigens in COVID-19 patients. Sci. Immunol. 5, (2020).

39. Elias, G. et al. Preexisting memory CD4 T cells in naïve individuals confer robust immunity upon hepatitis B vaccination. eLife 11, (2022).

40. Ionescu, L. & Urschel, S. Memory B Cells and Long-lived Plasma Cells. Transplantation 103, 890– 898 (2019).

41. Palm, A.-K.E. & Henry, C. Remembrance of Things Past: Long-Term B Cell Memory After Infection and Vaccination. Front. Immunol. 10, 1787 (2019).

42. Carmo, A. M. & Henriques, S. N. Cell activation and signaling in lymphocytes. in Tissue-Specific Cell Signaling (eds. Silva, J. V., Freitas, M.J. & Fardilha, M.) 133–161 (Springer International Publishing, 2020). doi:10.1007/978-3-030-44436-5_5.

43. Bourque, J. & Hawiger, D. Variegated outcomes of T cell activation by dendritic cells in the steady state. J. Immunol. 208, 539–547 (2022).

44. Zhang, J., Liang, Y. & Zhang, Y. Atomic-level protein structure refinement using fragment-guided molecular dynamics conformation sampling. Structure 19, 1784–1795 (2011).

45. Runthala, A. Refinement and improvement of template based protein modelling algorithms. (2015).

46. Heo, L., Park, H. & Seok, C. GalaxyRefine: Protein structure refinement driven by side-chain repacking. Nucleic Acids Res. 41, W384–8 (2013).

47. Dawson, D. V., Ozgur, M., Sari, K., Ghanayem, M. & Kostyu, D. D. Ramifications of HLA class I polymorphism and population genetics for vaccine development. Genetic Epidemiology (2001).

48. Bui, H.-H. et al. Predicting population coverage of T-cell epitope-based diagnostics and vaccines. BMC Bioinformatics 7, 153 (2006).

49. Vita, R. et al. The Immune Epitope Database (IEDB): 2018 update. Nucleic Acids Res. 47, D339– D343 (2019).

50. Wieczorek, M. et al. Major histocompatibility complex (MHC) class I and MHC class II proteins: conformational plasticity in antigen presentation. Front. Immunol. 8, 292 (2017).

51. Aiman, S. et al. Multi-epitope chimeric vaccine design against emerging Monkeypox virus via reverse vaccinology techniques-a bioinformatics and immunoinformatics approach. Front. Immunol. 13, 985450 (2022).

52. Tan, C., Zhu, F., Pan, P., Wu, A. & Li, C. Development of multi-epitope vaccines against the monkeypox virus based on envelope proteins using immunoinformatics approaches. Front. Immunol. 14, 1112816 (2023).

53. Wendorff, M. et al. Unbiased Characterization of Peptide-HLA Class II Interactions Based on Large-Scale Peptide Microarrays; Assessment of the Impact on HLA Class II Ligand and Epitope Prediction. Front. Immunol. 11, 1705 (2020).

54. Khare, S. et al. Gisaid’s role in pandemic response. China CDC Wkly 3, 1049–1051 (2021).

55. Aksamentov, I., Roemer, C., Hodcroft, E. & Neher, R. Nextclade: clade assignment, mutation calling and quality control for viral genomes. JOSS 6, 3773 (2021).

56. Katoh, K., Rozewicki, J. & Yamada, K. D. MAFFT online service: multiple sequence alignment, interactive sequence choice and visualization. Brief. Bioinformatics 20, bbx108 (2019).

57. Cock, P. J. A. et al. Biopython: freely available Python tools for computational molecular biology and bioinformatics. Bioinformatics 25, 1422–1423 (2009).

58. Zhang, K.-Y. et al. Vgas: A viral genome annotation system. Front. Microbiol. 10, 184 (2019).

59. McKinney, W. Data structures for statistical computing in python. in Proceedings of the 9th Python in Science Conference 56–61 (SciPy, 2010). doi:10.25080/Majora-92bf1922-00a.

60. Gonzalez-Galarza, F. F. et al. Allele frequency net database (AFND) 2020 update: gold-standard data classification, open access genotype data and new query tools. Nucleic Acids Res. 48, D783–D788 (2020).

61. Vita, R. et al. The immune epitope database (IEDB) 3.0. Nucleic Acids Res. 43, D405–12 (2015).

62. Doytchinova, I. A. & Flower, D. R. Bioinformatic approach for identifying parasite and fungal candidate subunit vaccines. Open Vaccine J. 1, 22–26 (2008).

63. Dimitrov, I., Bangov, I., Flower, D. R. & Doytchinova, I. AllerTOP v.2--a server for in silico prediction of allergens. J. Mol. Model. 20, 2278 (2014).

64. Gupta, S. et al. In silico approach for predicting toxicity of peptides and proteins. PLoS ONE 8, e73957 (2013).

65. Gasteiger, E. et al. Protein identification and analysis tools on the expasy server. in The proteomics protocols handbook (ed. Walker, J. M.) 571–607 (Humana Press, 2005). doi:10.1385/1-59259-890-0:571.

66. Altschul, S. F., Gish, W., Miller, W., Myers, E. W. & Lipman, D. J. Basic local alignment search tool. J. Mol. Biol. 215, 403–410 (1990).

67. Yang, J. et al. The I-TASSER Suite: protein structure and function prediction. Nat. Methods 12, 7–8 (2015).

68. McGuffin, L. J., Bryson, K. & Jones, D. T. The PSIPRED protein structure prediction server. Bioinformatics 16, 404–405 (2000).

69. Williams, C. J. et al. MolProbity: more and better reference data for improved all-atom structure validation. Protein Sci. 27, 293–315 (2018).

70. Schrödinger, L. The PyMOL Molecular Graphics System, Version 1.8. (No Title) (2015).

71. Rapin, N., Lund, O., Bernaschi, M. & Castiglione, F. Computational immunology meets bioinformatics: the use of prediction tools for molecular binding in the simulation of the immune system. PLoS ONE 5, e9862 (2010).

72. Binette, V., Mousseau, N. & Tuffery, P. A Generalized Attraction-Repulsion Potential and Revisited Fragment Library Improves PEP-FOLD Peptide Structure Prediction. J. Chem. Theory Comput. 18, 2720–2736 (2022).

73. Rey, J., Murail, S., de Vries, S., Derreumaux, P. & Tuffery, P. PEP-FOLD4: a pH-dependent force field for peptide structure prediction in aqueous solution. Nucleic Acids Res. 51, W432–W437 (2023).

74. Chen, F. et al. Assessing the performance of the MM/PBSA and MM/GBSA methods. 6. Capability to predict protein-protein binding free energies and re-rank binding poses generated by protein-protein docking. Phys. Chem. Chem. Phys. 18, 22129–22139 (2016).

75. Feng, T. et al. HawkRank: a new scoring function for protein-protein docking based on weighted energy terms. J. Cheminform. 9, 66 (2017).

76. Velankar, S. et al. Pdbe: protein data bank in europe. Nucleic Acids Res. 38, D308–17 (2010).

